# Efficient high-resolution TMS mapping of the human motor cortex by nonlinear regression

**DOI:** 10.1101/2021.03.11.434996

**Authors:** Ole Numssen, Anna-Leah Zier, Axel Thielscher, Gesa Hartwigsen, Thomas R. Knösche, Konstantin Weise

**Author notes:** CORRESPONDING AUTHOR, Ole Numssen; Max Planck Institute for Human Cognitive and Brain Sciences, Stephanstr. 1a, 04103 Leipzig, Germany.

## Abstract

Transcranial magnetic stimulation (TMS) is a powerful tool to investigate causal structure-function relationships in the human brain. However, a precise delineation of the effectively stimulated neuronal populations is notoriously impeded by the widespread and complex distribution of the induced electric field.

Here, we propose a method that allows rapid and feasible cortical localization at the individual subject level. The functional relationship between electric field and behavioral effect is quantified by combining experimental data with numerically modelled fields to identify the cortical origin of the modulated effect. Motor evoked potentials (MEPs) from three finger muscles were recorded for a set of random stimulations around the primary motor area. All induced electric fields were nonlinearly regressed against the elicited MEPs to identify their cortical origin.

We could distinguish cortical muscle representation with high spatial resolution and localized them primarily on the crowns and rims of the precentral gyrus. A post-hoc analysis revealed exponential convergence of the method with the number of stimulations, yielding a minimum of about 180 random stimulations to obtain stable results.

Establishing a functional link between the modulated effect and the underlying mode of action, the induced electric field, is a fundamental step to fully exploit the potential of TMS. In contrast to previous approaches, the presented protocol is particularly easy to implement, fast to apply, and very robust due to the random coil positioning and therefore is suitable for practical and clinical applications.

## Introduction

Transcranial magnetic stimulation (TMS) is a powerful non-invasive technique to modulate motor and cognitive functions in the human brain through inducing electric fields. This provides insight into structure-function relationships at both group and individual level (e.g. [1,2]). However, the precise cortical location at which an induced electric field initiates the behavioral effect remains unclear. Estimating the induced fields is key to address this [1,2] but complex field distributions and interindividual variations hamper precise cortical localization. For example, the regions which are initially activated for TMS-elicited motor evoked potentials (MEP) are still debated. Several studies point to crown and lip stimulation in the caudal part of the dorsal premotor cortex (PMd, situated in BA6d) [1,3–5]. Others point towards a direct stimulation of the hand representations [6,7] in the primary motor cortex (M1, situated in BA4a/BA4p), inside the sulcal wall of the precentral gyrus (Fig. 1). Localization efforts are considerably impeded by substantial inter-individual variation of the cortical geometry [8–13].

**Figure 1:**
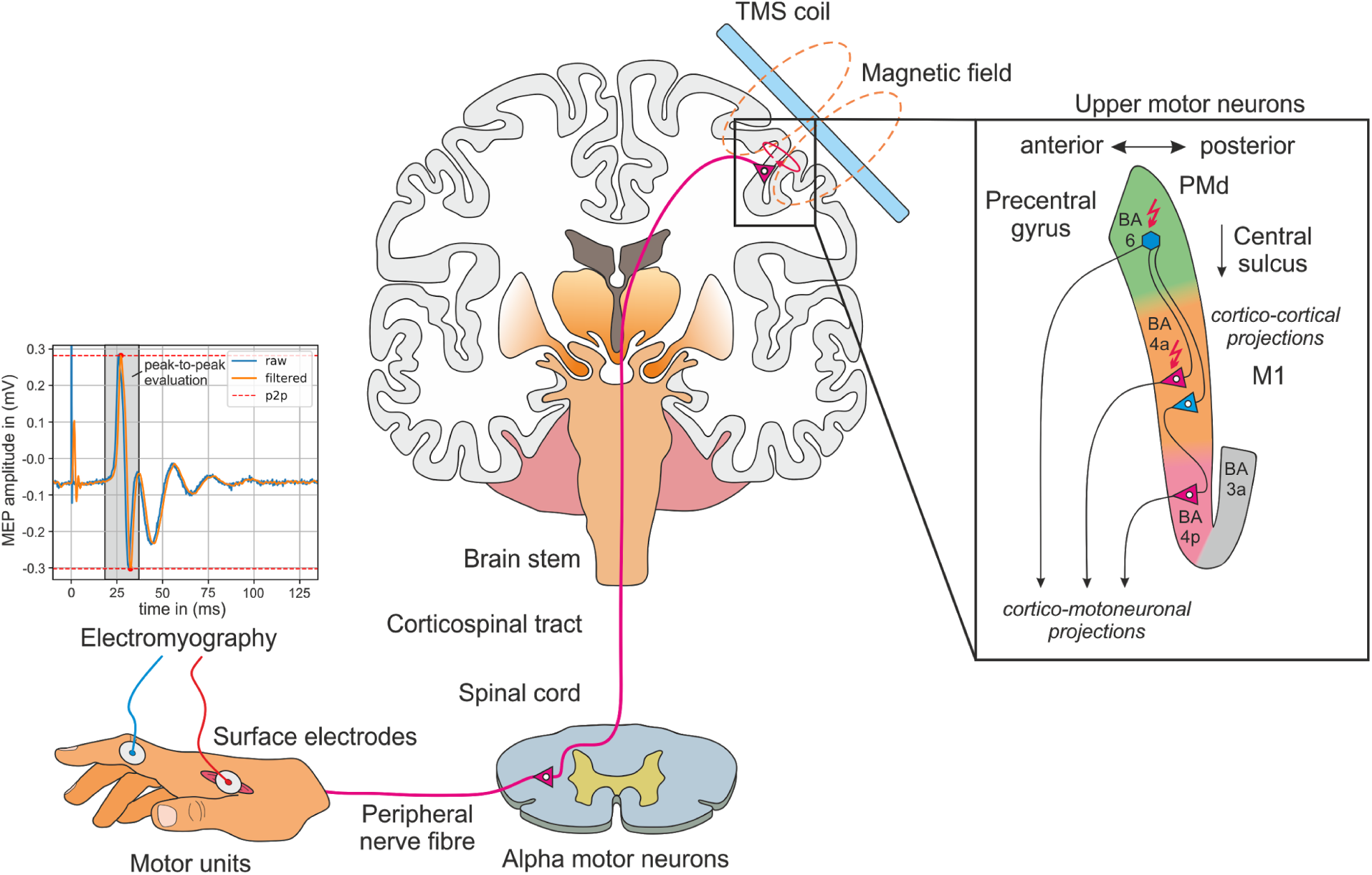
Schematic representation of TMS experiments measuring motor evoked potentials (MEPs) from finger muscles. The TMS coil is located tangentially to the scalp over the primary motor cortex. A time varying current in the coil generates a time varying magnetic field, which induces an electric field in the brain. It is still debated which regions are initially activated when motor evoked potentials (MEP) are elicited by TMS. Candidate mechanisms include: (i) direct stimulation of pyramidal output neurons in the primary motor cortex (M1, area BA4a/p); (ii) indirect depolarization of M1 output neurons via cortico-cortical projections from the dorsal premotor cortex (PMd, part of BA6d) to M1; or (iii) stimulation of pyramidal output neurons in PMd and direct cortico-motoneuronal projections to alpha motor neurons. Summed muscle activation is quantified via electromyography. Upper motor neuron details adapted from [25].

Many previous mapping studies in the motor cortex (e.g. [14–18]) rely on center of gravity approaches and structured grids. This allows for the identification of those TMS coil positions on the head surface that yield the strongest effect. Frameworks have been proposed to reduce user-dependence [19], relying on custom built TMS hardware to identify the optimal coil position in an automated way. These coil configurations can then be projected onto the cortex to estimate functionally relevant brain structures in a simplified manner [7,20]. However, these approaches can only provide a rough estimate of the effective stimulation site because information about the induced electric field is disregarded. Furthermore, no depth information is included and potential spatial offsets between the electric field peaks from a position directly under the coil are neglected.

Recently, localization methods have been improved by considering the interaction between estimated electric fields and observed effects, for example by superimposing multiple fields in an additive or multiplicative fashion [21,22]. Yet, such approaches suffer from the fundamental problem that the field maximum cannot be positioned freely throughout the cortex. Brain areas close to the TMS coil always receive higher field strengths than deeper located areas, leading to a strong bias in favor of superficial regions. To overcome this, Bungert et al. [23] employed a statistical approach based on the resting motor threshold (rMT) for different coil orientations. Others [24] investigated the influence of the coil position. We recently introduced an approach exploiting information from both coil positions and orientations [1]. This linked the induced electric field strength to a measure quantifying the TMS effect (e.g., MEP amplitude). This relationship, the input-output curve (IO-curve), is evaluated at every cortical element of a fine mesh for a set of different coil positions and orientations. In contrast to previous studies, the approach was validated in an additional TMS experiment by explicitly testing whether stimulation with an optimized coil position and orientation for the proposed cortical locations yielded a larger effect than stimulation with deviating coil positions or orientations. This method was robust against measurement and tissue conductivity uncertainties. With only six optimal coil positions and orientations, resulting in about 600 TMS pulses, the localization problem could be solved. However, it remained unclear how to identify these positions and orientations. Here, we substantially advanced this method to allow arbitrary (‘random’) coil positions and orientations for each TMS pulse and thus providing a simple experimental protocol. This decreases the required number of stimulations for reliable localization, by virtue of increased electric field variability. We applied this method to 13 subjects to identify the somatotopic organization of three hand muscles within one TMS experiment. The proposed cortical muscle representations were validated in a second experimental session to show successful distinction of different finger muscle representations at the individual subject level. Based on an extensive convergence analysis, we provide metrics to evaluate the overall goodness of the proposed cortical mapping procedure.

## Materials and methods

### Subjects

Thirteen healthy, right-handed participants (six females, age 21-38 years) with a mean laterality index of 94.92 (*SD* = 8.13) according to the Edinburgh Handedness Inventory were recruited. Subject inclusion was in accordance with the safety guidelines for TMS studies [26]. Written informed consent was obtained from all participants prior to the examination. The study was performed according to the guidelines of the Declaration of Helsinki and approved by the local Ethics committee of the Medical Faculty of the University of Leipzig.

### Hardware setup

TMS pulses were applied with a MagPro X100 stimulator (MagVenture, firmware Version 7.1.1) and a MCF-B65 figure-of-eight coil. Coil positioning was guided by a neuronavigation system (TMS Navigator, Localite, Germany, Sankt Augustin; camera: Polaris Spectra, NDI, Canada, Waterloo) and the coil positions were saved for each stimulation.

Electromyographic data (EMG) for three hand muscles were recorded from the subjects’ right hand for each stimulation, namely the first dorsal *interosseous* (FDI), the musculus abductor *digiti minimi* (ADM), and the musculus abductor *pollicis brevis* (APB) using a standard belly-tendon montage [15]. Electrodes were connected to a patient amplifier system (D-360, DigitimerLtd., UK, Welwyn Garden City; bandpass filtered from 10 Hz to 2 kHz), which was connected to an acquisition interface (Power1401 MK-II, CED Ltd., UK, Cambridge, 4 kHz sampling rate). EMG recording was performed with Signal (CED Ltd., version 4.11). Subsequently, EMG data was lowpass filtered with a 6^th^ order Butterworth filter (cutoff frequency: 500 Hz). MEPs were calculated as peak-to-peak amplitudes in a time window 18 to 35 ms after the TMS pulse.

### Localization experiment

Localization of the initial MEP target location was initially guided by group coordinates from [27]. We manually determined the rMT for the FDI and the corresponding coil position [28]. The rMT was defined as the minimum stimulator intensity yielding MEPs with an amplitude of at least 50 μV in at least 5 out of 10 consecutive TMS pulses [29].

900-1100 single biphasic pulses were applied with a 5 s inter stimulus interval. Coil positions and angles were randomly selected for each stimulation to sample electric field distributions and corresponding MEPs for different induced electric fields. The coil center positions were restricted to a circular area of 2 cm radius around the estimated M1 location (Fig. 2) and the angles to approximately ±60° in relation to the estimated optimal coil angle. The stimulation area definition proved sufficient to yield MEPs amplitudes from the upper limit of the I/O curve at the estimated M1 stimulation site and small or no MEPs at the periphery for a fixed stimulator intensity. This intensity, about 150% of the estimated rMT, was individually predetermined to stay in a comfortable range. Experimenters were instructed to evenly sample the determined area.

**Figure 2.**
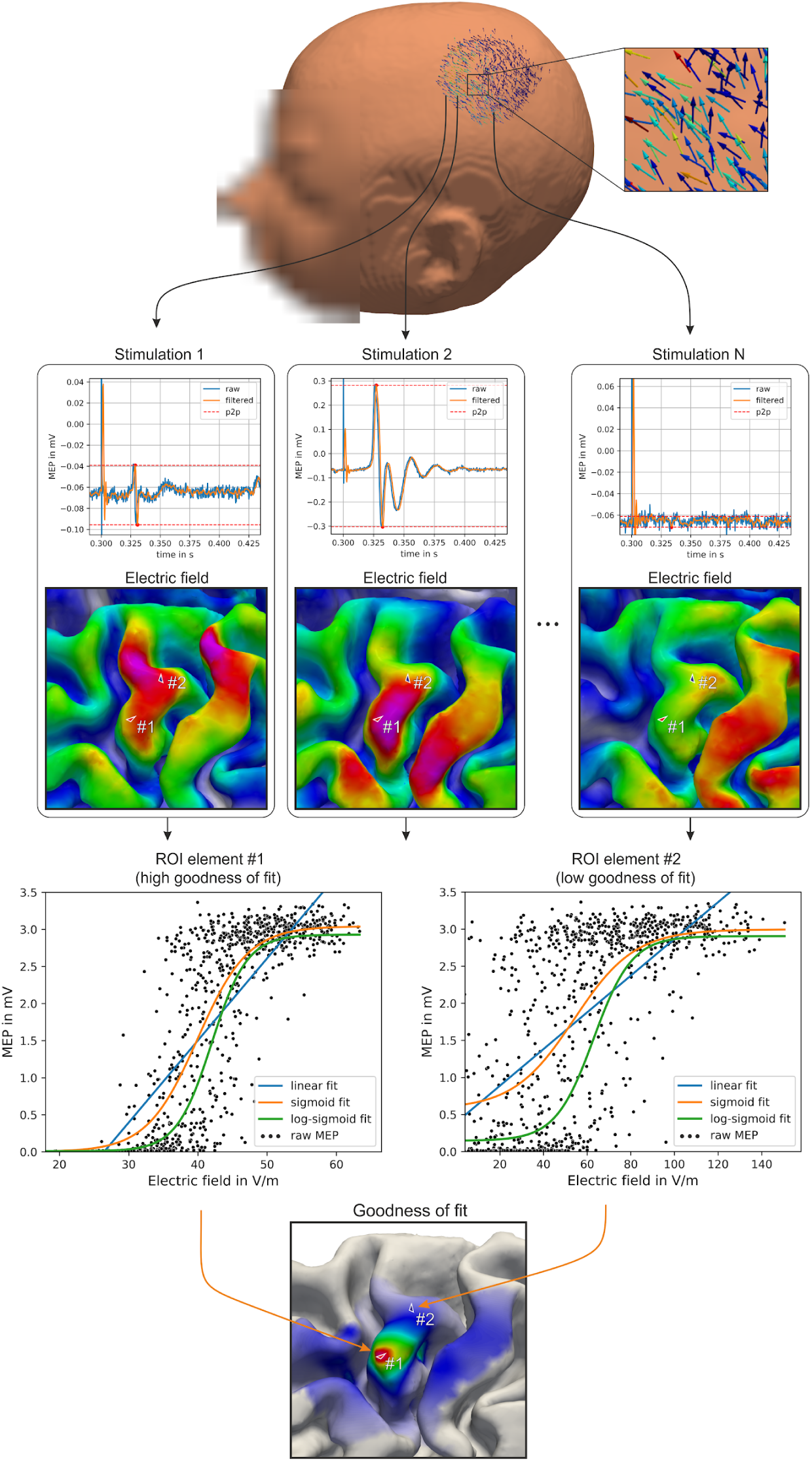
General principle of the localization approach. Top: *N* TMS pulses are applied at random coil locations and orientations around the M1 area. Stimulator intensity is calibrated such that large MEPs are elicited at the center of the examined area. For each pulse, a motor evoked potential (MEP) is recorded with muscle EMG and the induced electric field is calculated. Middle: For each cortical element, the electrical field quantity of interest (e.g. its magnitude) is regressed on the MEP amplitude. Bottom: A goodness-of-fit map identifies the most probable origin of the MEPs.

### 2.4 Numerical simulations of the induced electric field

Electric field calculations were conducted for each pulse (SimNIBS v3.1, [30,31]) with high-resolution anisotropic finite element models (FEMs). Magnetic resonance imaging (MRI) scans were acquired on a 3T MRI scanner (Siemens Verio or Skyra) with a 32 channel head coil using the same acquisition parameters as before [1]. T1 and T2 images were used for tissue type segmentation. Conductivity tensors in grey and white matter were reconstructed from diffusion weighted images using the volume normalized mapping approach (dwi2cond, https://simnibs.github.io/simnibs/build/html/documentation/command_line/dwi2cond.html, [32]). Individual head models were generated using the headreco pipeline [33] utilizing SPM12 (https://www.fil.ion.ucl.ac.uk/spm/software/spm12/, [34]) and CAT12 (http://www.neuro.uni-jena.de/cat/, [35]). The final head models were composed of ~3.4·10^6^ nodes and ~18.5·10^6^ tetrahedra (average volume: ~0.15 mm^3^ in the cortex). Six tissue types were included with the following conductivity estimates: white matter (*σ_WM_* = 0.126 *S/m*), grey matter (*σ_GM_* = 0.275 *S/m*), cerebrospinal fluid (*σ_CSF_* = 1. 654 *S/m*), bone (*σ_B_* = 0.01 *S/m*), skin (*σ_S_* = 0.465 *S/m*), and eyeballs (*σ_EB_* = 0.5 *S/m*) [31,36]. See [30] for FEM details.

A region of interest (ROI) was defined around the handknob area (FreeSurfer, (http://surfer.nmr.mgh.harvard.edu/, [37,38]) based on the fsaverage template. This covered parts of somatosensory cortex (BA1, BA3), primary motor cortex M1 (BA4), and dorsal premotor cortex PMd (BA6). The HCP-MMP parcellation was used [39] to visualize the border between PMd and M1. All analyses were performed on the grey matter midlayer (halfway between grey and white matter surfaces).

### Determining the site of effective stimulation

The core concept of the proposed method is illustrated in Fig. 2. We assume i) the existence of a sigmoidal relationship between electric field and MEP amplitude at the cortical location where the electric field elicits the observed MEP; ii) that this relationship is independent of the origin of the field, i.e. the coil position and orientation it was generated with; iii) that for each muscle exactly one focal cortical area in the primary motor cortex is functionally relevant for MEPs evocation. Exploiting these assumptions, the site of effective stimulation can be pinpointed by fitting cortical I/O curves [1], relating the local electric field to the MEP amplitude, and subsequently quantifying their goodness-of-fit. The goodness-of-fit would be highest at the cortical site that houses the relevant neuronal populations. Fitting can be performed for different components of the electric field vector, such as its magnitude (|*E*|), its projection onto the cortical surface normal (|*E*_⊥_|), its projection into the cortical tangential plane (|*E*_∥_|), and any other quantities derived thereof. Based on findings from experimental [1] and modeling approaches [40], we focus on the field magnitude. Fitting can be performed with different functions. We compared (i) standard linear regression, (ii) nonlinear regression using a sigmoidal function, and (iii) nonlinear regression using a log-transformed sigmoidal function.

(i) In standard linear regression, a linear relationship between electric field intensity *x_ij_* of stimulation i at the cortical element j and the MEP amplitude *y_i_* is assumed:

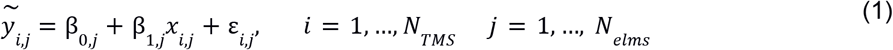

where *N_TMS_* and *N_elms_* denote the total number of applied TMS pulses and number of elements in the examined region of interest (ROI) respectively. β_0_ and β_1_ are estimated in every ROI element *j*. This approach is the cheapest in terms of computational cost but neglects the characteristic sigmoidal shape of the input-output curve observed in the motor cortex.
(ii) A sigmoidal input-output function provides more physiological plausibility:

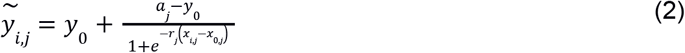

where *a* is the saturation amplitude, *r* is the slope, *X*_0_ is the location of the turning point on the abscissa, and *y*_0_ denotes the offset. The offset parameter *y*_0_ results from measurement noise. We estimated *y*_0_ from baseline EMG data in the absence of any stimulus [41] to reduce the number of free parameters.
(iii) EMG data from this domain are typically heteroscedastic and log-transformation may be applied to equalize variance across the range of MEP magnitudes [42–45]. Here, we used a sigmoidal function of the following type:

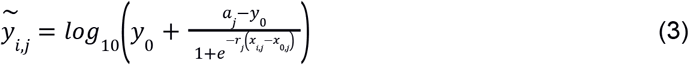 The non-linear functions were fitted in every ROI element using the Levenberg-Marquardt algorithm. We assessed the element-wise goodness-of-fit by means of the coefficient of determination *R*^2^:

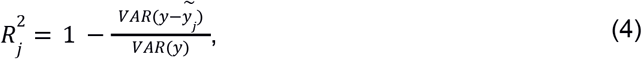

where *y* and 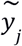 are vectors containing measured and fitted MEPs, respectively. The better the fit, the closer *R*^2^ is to unity.

The proposed method is principally equivalent to the curve shift approach proposed previously [1]. However, in our approach, curve fitting is done element-wise in the E-MEP space leveraging information from every single TMS pulse, in contrast to the condition-wise approach implemented before. The procedure yields one cortical map of *R*^2^ scores per muscle, quantifying the probability of generating MEPs for each cortical element.

### Validation experiment

After identifying the neuronal population that appears to underlie the observed MEP, i.e. yielding the highest *R*^2^ score, we validated this finding in a second experimental session. We determined, for each of the three muscle representations, the optimal coil position and orientation to stimulate the identified cortical location (‘hotspot’). To verify if stimulation of the proposed hotspot does indeed lead to the largest effect, rMTs were acquired for the optimal coil positions/orientations and for adjacent ones. Optimal coil positions/orientations were determined with an extensive search procedure, comparing electrical fields magnitudes of 4852 coil configurations at the hotspot area (search radius = 20 mm, spatial resolution = 2.5 mm, search angle = 180°, angle resolution = 7.5°). The optimization routine is implemented in SimNIBS v3.1 (https://simnibs.github.io/simnibs) and has been described before ([1]; Supplementary Material). To determine the true rMT, single biphasic pulses with 5 s interstimulus intervals were applied. The same figure-of-eight coil as before was used here. The rMTs for optimal coil configurations were compared to rMTs obtained for six adjacent coil configurations. Two coil configurations shared the same coil center with the optimal one but differed in their coil orientation (−45° and +45°) and four coil configurations shared the optimal coil orientation, but were shifted 7.5 mm into superior, posterior, inferior, or anterior directions.

### Convergence analysis

To identify the minimum number of pulses needed to carry out the cortical localization, we studied random subsets of the realized stimulations, from n = 10 to all available (900-1100) stimulations. We drew 100 independent subsets for each subject to approximate the robustness.

We assessed two metrics: i) the normalized root mean square deviation (NRMSD) as a measure of overall shape similarity of the resulting cortical map and ii) the geodesic distance to quantify an estimate of the accuracy of the hotspot identification. Convergence for both was quantified against the full set of stimulations as well as against the previous solution from *n – 1* stimulations. The former yields a proxy for the (unknown) ground truth, whereas the latter quantifies the magnitude of change from one stimulation to the next. This stability measure can be used in online analyses to construct a stop criterion, as well as for post-hoc evaluation of the overall goodness of the mapping procedure.

The NRMSD between the 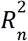 map for *n* stimulations and the reference map 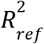 was calculated as:

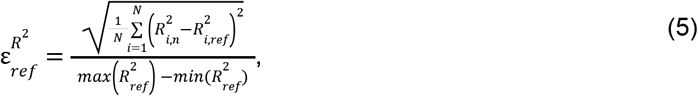

The geodesic distance 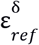 was calculated with tvb-gdist 2.1.0 (github.com/the-virtual-brain/tvb-gdist). To estimate the generalizability of our results, we carried out non-parametric permutations tests across the stimulation subsets. These were performed against the full set solution. Based on this, the lower bound of the number of stimulations, which was needed to reach < 5% error NRMSD, could be derived. For the geodesic distance a criterion of < 5 mm was chosen.

## Results

### Localization

For each subject, we calculated the cortical mapping with linear, sigmoidal, and log-transformed sigmoidal functions to identify the representations of APB, FDI, and ADM. Detailed results are presented in Fig. 3 for one subject considering the electric field magnitude (see Fig. S1 and S2 for the tangential and normal component, respectively). The general shape of the normalized *R*^2^ maps are very similar across different fitting functions.

**Figure 3:**
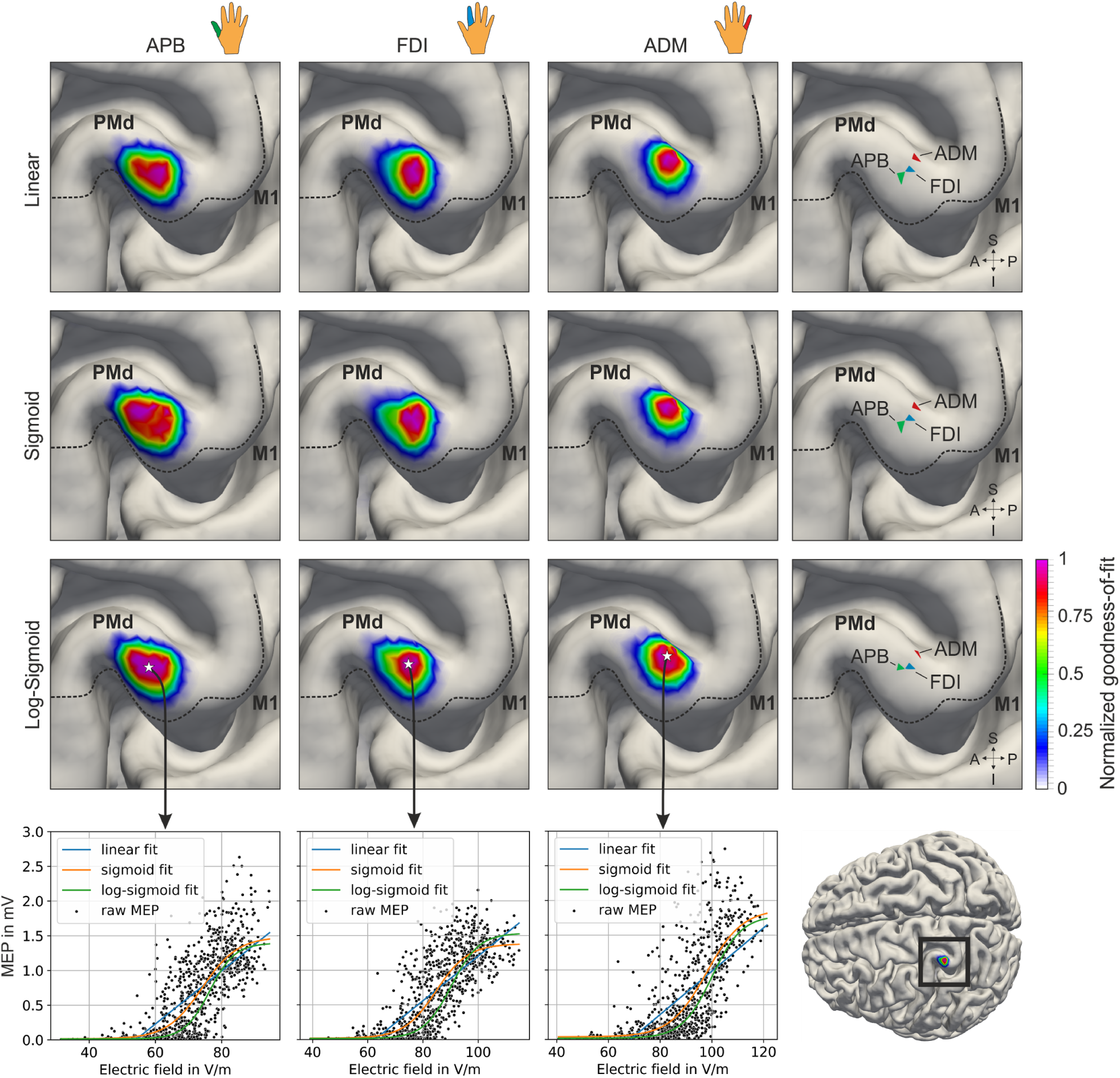
Log-sigmoidal fitting exhibits an optimal yield-cost tradeoff. The first three columns show the normalized coefficients of determination (*R*^2^) for APB, FDI, and ADM, respectively, considering the magnitude of the electric field for subject S12. Absolute values range between 0.3 for linear regression and 0.7 for non-linear regression. The last column highlights the identified hotspots for the three digits. Mapping results for linear, sigmoidal and log-transformed sigmoidal functions are of a similar shape (rows). The boundary between dorsal premotor cortex PMd/BA6 and the primary motor cortex M1/BA4 are determined with the HCP-MMP atlas [39]. Function fits for optimal elements are characterized by a clear relation between the induced electric field strength and the evoked motor potentials (bottom). *R*^2^ values were potentiated by an exponent of 20 and scaled to [0,1] range per muscle and fit function for visualization purposes.

The computation time for sigmoidal mappings was significantly shorter than for mappings with log-transformed sigmoids. Thus, we used sigmoidal functions throughout the rest of the analyses as this function type provides a well-balanced compromise in terms of mapping accuracy and computation efficacy. Table S1 provides peak lvalues for all methods per subject.

Mapping results for all 13 subjects considering the electric field magnitude are shown in Fig. 4 (see Fig. S3 and S4 for the tangential and normal component, respectively). The *R*^2^ hotspots are primarily located on the gyral crown and rim of the precentral gyrus. Representations of FDI and APB were found to be located closer to each other than to ADM, which was generally situated superior to them. For one subject, the muscle representations were located in M1, whereas for 12 subjects, they were located in caudal PMd. See Table 1 for group average results.

**Figure 4:**
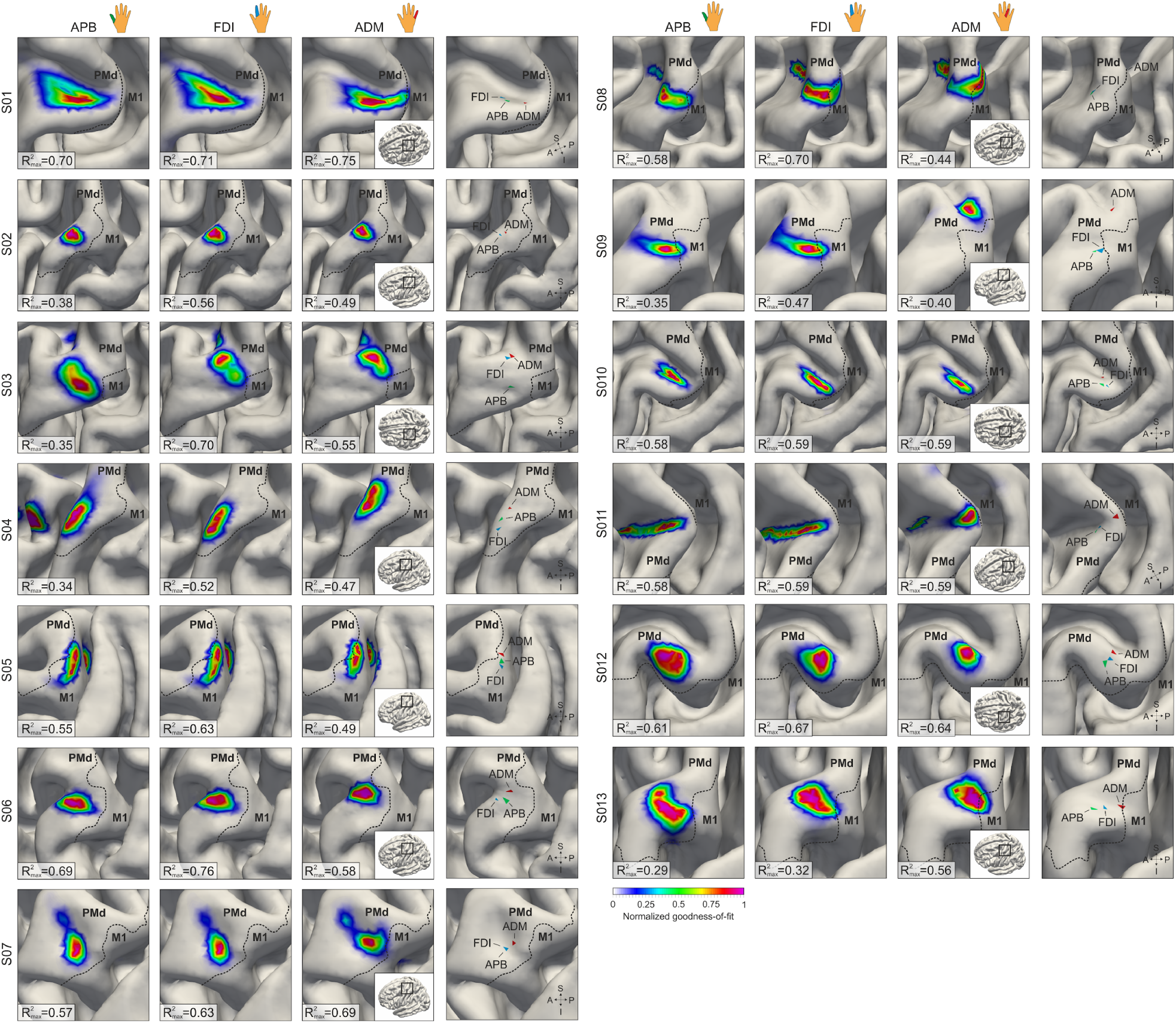
Localization results from all 13 subjects using nonlinear regression with a sigmoidal function. First three columns: normalized coefficients of determination (*R*^2^) for APB, FDI, and ADM, respectively, considering the magnitude of the electric field. Absolute values range between 0.4 and 0.8. The last column highlights the identified hotspots for the three digits. Subject S01 was measured with random stimulation intensities, instead of the fixed intensity that was used otherwise. The boundary between dorsal premotor cortex PMd/BA6 and the primary motor cortex M1/BA4 was determined with the HCP-MMP atlas [39]. *R*^2^ values were potentiated by an exponent of 20 and scaled to [0,1] range per muscle and fit function for visualization purposes.

**Table 1:**
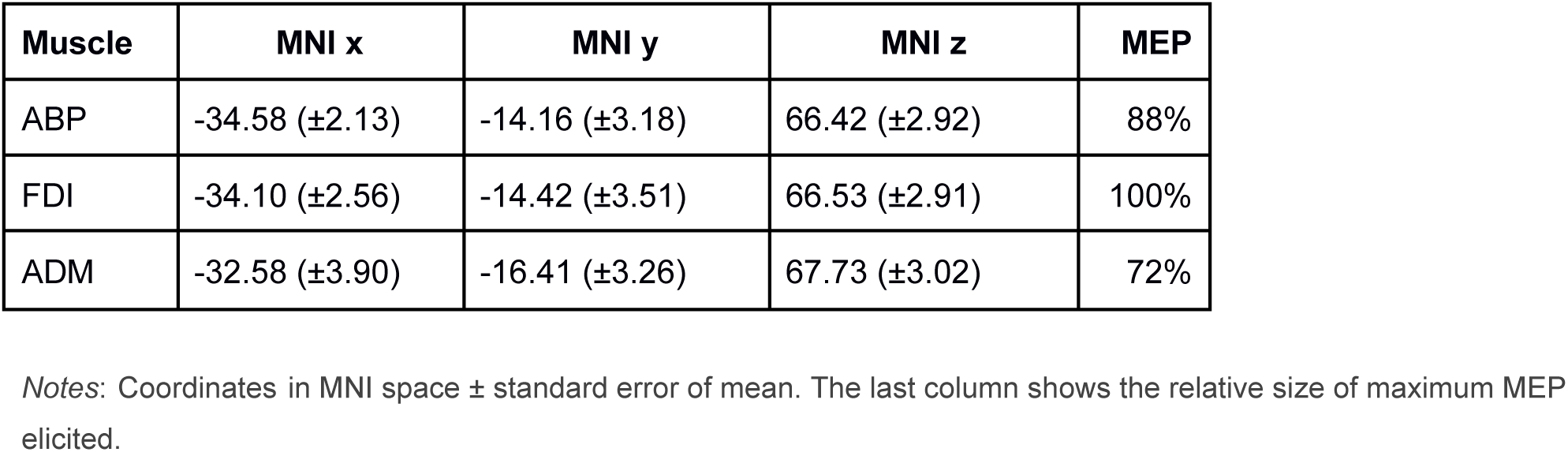
Group-average results for the three finger muscles.

Localization considering the magnitude and the tangential component yielded similar results, whereas no clear relationship was found for the normal component.

### Validation

After identifying the cortical digit hotspots, we determined optimal coil positions to stimulate these hotspots and measured rMTs for these and for adjacent coil configurations. to validate the mapping results. Of the 13 initial subjects, eight participated in the validation study (Fig. 5). In the majority of cases the rMT for the optimal coil position was the lowest. Because data was not distributed normally (Shapiro-Wilk test: *W* = 0.94215, *p* = 2.432*e* - 06) one Mann-Whitney test was performed for each muscle. Resting MTs at adjacent positions were significantly higher than rMTs from the optimal, pre-computed coil positions and orientations (APB: *W* = 316, *p* = 1.709*e* - 03; FDI: *W* = 332, *p* = 4.755*e* - 04; *W* = 356, *p* = 5. 784*e* - 05).

**Figure 5:**
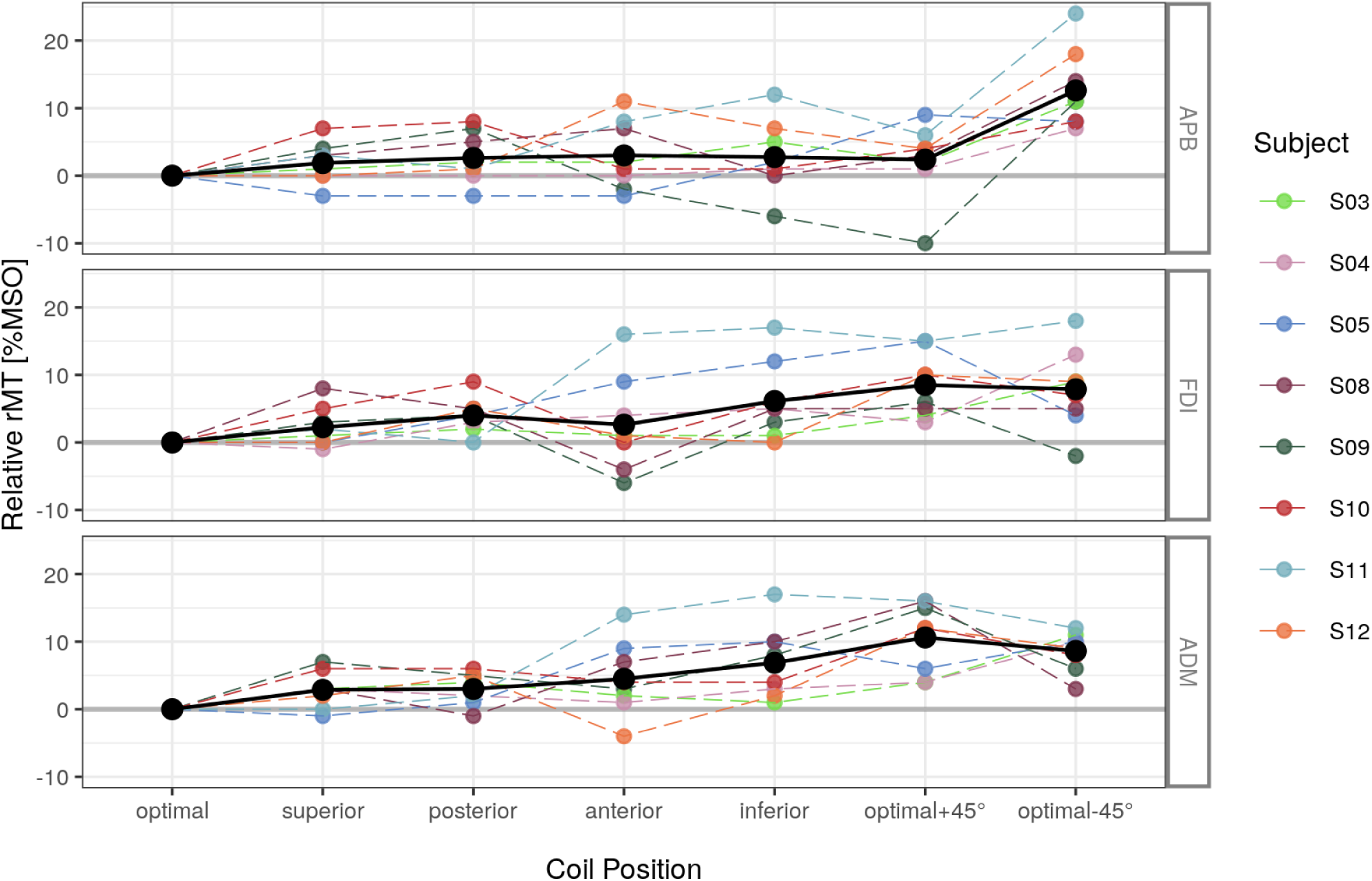
Pre-computed optimal coil positions produce minimum resting motor thresholds (rMTs). Six validation positions were tested: four positions with same optimal coil rotation: *superior, posterior, anterior, inferior*; two coil orientations at the optimal coil center position: *optimal −45°, optimal +45°*. Resting motor thresholds for each were assessed and normalized to the rMT of the pre-computed optimal position per subject and muscle. Color: subject id. Black: average rMT %MSO: Percentage of maximum stimulator output. APB: thumb. FDI: index finger. ADM: little finger. Explicit rMT values are provided in the supplemental material Table S2.

### Convergence analysis

For 100 random sequences, we calculated the mapping ranging from *n* = 10 to all available stimulations (*N*). NRMSD and geodesic distance converged exponentially, tested against all available stimulations *(N)* and against the previous solution (*n* - 1). The variance across these localizations was large only until about *n* = 50 and then quickly decreased. The convergence properties are comparable between subjects for the NRMSD metric (Fig. 6), and distinctively more pronounced for the geodesic distance (Fig. 7). Across subjects and metrics, localizing FDI was feasible with fewer stimulations than localizing ADM and APB.

**Figure 6:**
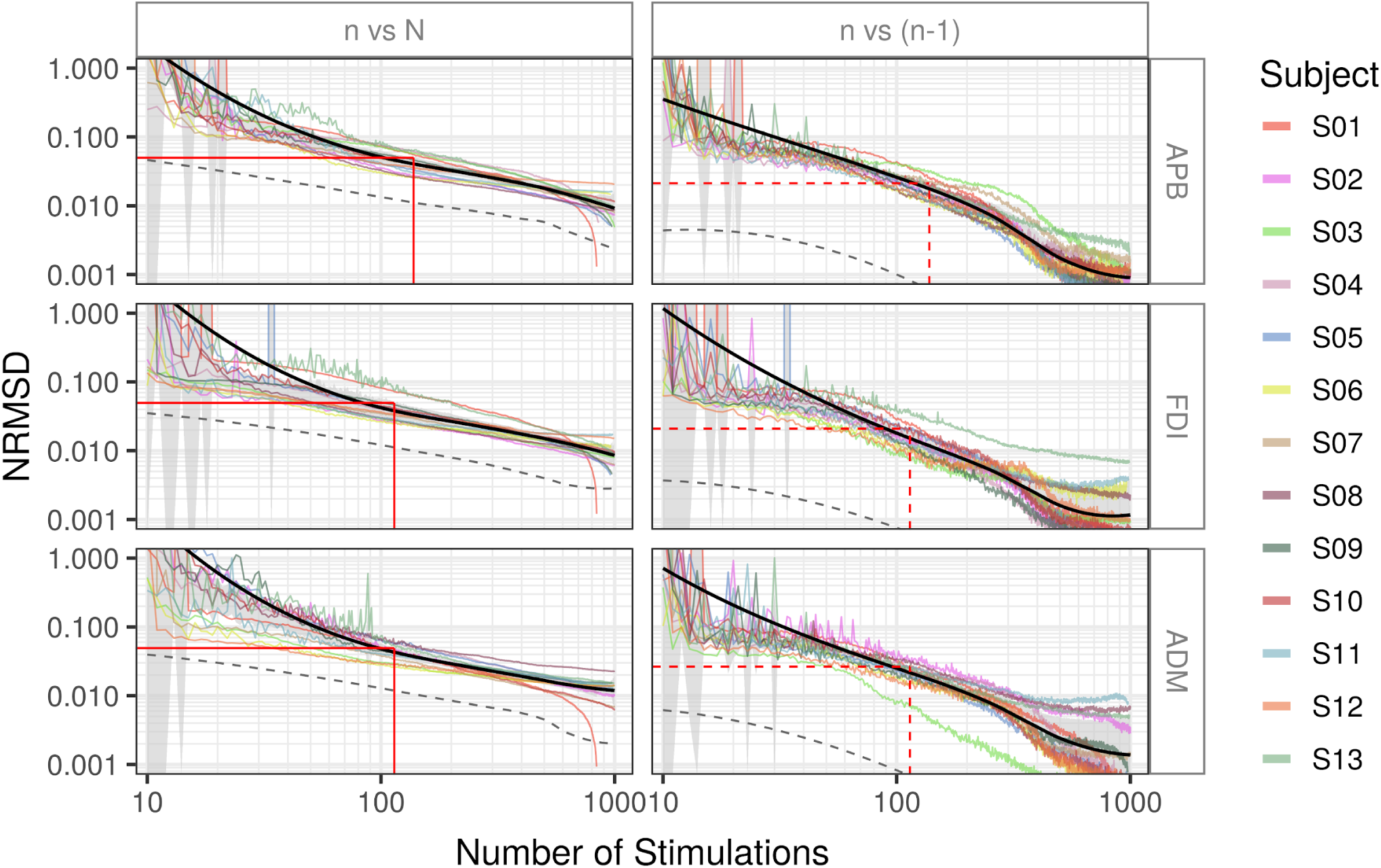
Localization maps converge after approximate 150 random stimulations. Normalized root mean square deviation (NRMSD), a proxy of overall similarity of solutions, as a function of *n* randomly selected stimulations. Left: NRMSD against the reference solution from all:*N*≈1000 stimulations. Right: NRMSD against the previous *n* − 1 stimulations. Colored lines: subject-wise average convergence across 100 random samples (see text). Black line: grand average, smoothed for the sake of visualization. Grey area: confidence interval of population mean based on nonparametric bootstrapping at α = 0.95 level. Grey dashed line: best solution across samples. The number of stimulations needed to reach 5% NRMSD against the reference solution *N* is depicted left (red solid lines). This number of stimulations and the corresponding NRMSD value for the *n* − 1 comparison is delineated with red dashed lines for the *n* − 1 comparison (right).

**Figure 7:**
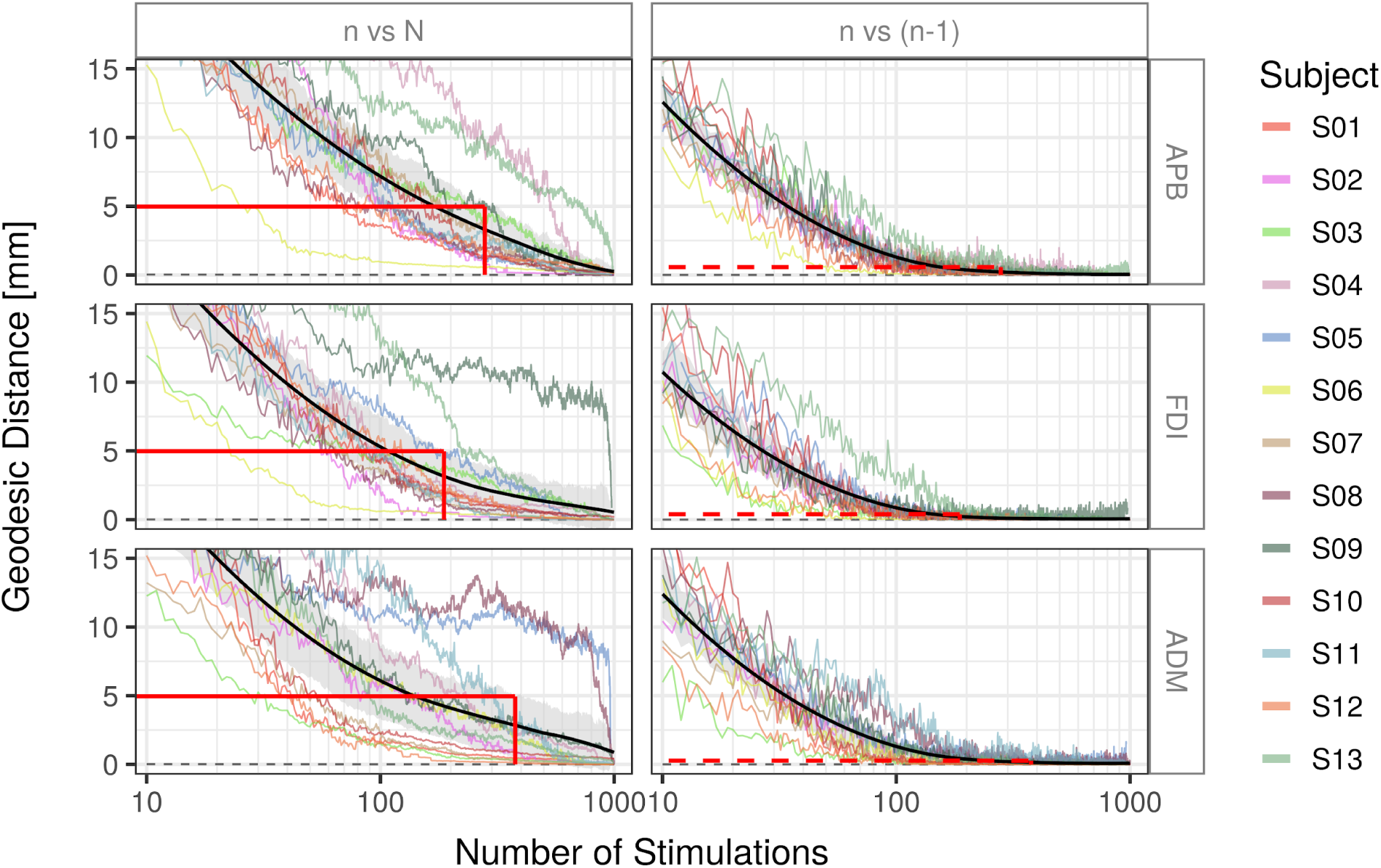
Robust identification of the correct target element. Geodesic distance (in mm) of the identified target from *n* stimulations to the target identified by the reference solution. Left: reference solution is based on all *N*≈1000 stimulations. Right: reference is the previous solutions from *n* − 1 stimulations. Colored lines: subject-wise average convergence (see text). Black line: grand average. Grey area: confidence interval around population mean based on nonparametric bootstrapping at *α* = 0.95 level. Grey dashed line: best solution across samples. The number of stimulations needed to reach 5 mm distance against the reference solution *N* is depicted left (red solid line). This number of stimulations and the corresponding distance is delineated with a dashed red line for the *n* − 1 comparison (right).

With non-parametric bootstrapping tests we identified the number of stimulations required to localize within a 95% confidence interval. Tests were performed for the comparison against all stimulations as a proxy for the ground truth. 138, 114, and 114 stimulations are necessary for APB, FDI, and ADM, respectively, for the overall similarity of *R*^2^ maps (NRMSD) to reach an error <5%. This corresponds to NRMSD values of 2.1%, 2.0%, and 2.6% (APB, FDI, ADM) for comparisons against the previous solution (*n* - 1). Correct cortical targets can be pinpointed within 5 mm with 280, 187, and 377 (APB, FDI, ADM) stimulations as quantified by the geodesic distance. This corresponds to 0.57 mm, 0.39 mm, and 0.26 mm distance when comparing against the previous solution (*n* - 1).

See Vid. 1 for a video demonstration of the FDI *R*^2^ map convergence behavior for a representative subject.

## Discussion

We propose an efficient and high-resolution method to identify structure-function relationships with TMS and applied this method to map the individual somatotopy of several hand muscle representations in the primary motor cortex. This approach links information about the induced electric fields at the cortex to a modulated experimental outcome and relies on the variance across induced electric fields from multiple different stimulation sites.. It is easy to implement in a standard TMS laboratory because the localization does not depend on predetermined sensitive coil positions and orientations and is therefore more suitable for practical and clinical use than previously proposed mapping approaches.

Specifically, we applied TMS pulses with randomly chosen coil positions and orientations around the larger primary motor region and recorded motor evoked potentials (MEPs) from several hand muscles. Subsequent electric field modelling and biophysical informed analyses allowed us to identify the associations between electric field magnitude and MEP size in the cortex. In line with previous work [1], hotspots were found on the crowns and rims of the precentral gyrus between BA6 and BA4 for the magnitude and the tangential component of the electric field, while no clear relationship was found for the normal component.

Based on our results, we reason that TMS either evokes muscle activity through direct cortico-motoneuronal pyramidal neurons from the caudal part of PMd (PMdc) [46] or indirectly via cortico-cortical premotor-motor (PMdc-to-BA4a/p) projections [47]. This is supported by studies reporting that TMS during periods of high surface negativity measured by EEG leads to higher MEP amplitudes [48-50]. This surface negativity may specifically stem from apical dendrites of neurons with a radial cortical orientation [51], which are located in the crown of the precentral gyrus. Since the exact relationship between oscillations and corticospinal excitability remains unknown, future work on accurate neuronal network models is necessary to address this question.

The observed geodesic distances between finger representations are consistent with recent high-resolution functional MRI studies [52]. Moreover, the general somatotopic organization of hand muscle representations in the primary motor cortex is in accordance with previous work, with APB being located infero-lateral to ADM [53,54] and ADM superior to FDI [5,55-57]. The observed substantial variability between subjects is also consistent with previous studies [58].

We extensively assessed the convergence behavior of the presented mapping method. Thereby, we were able estimate a lower bound for the number of random stimulations needed to achieve robust somatotopic maps. About 150 pulses from random coil positions and orientations suffice to obtain robust cortical maps. This yields a mapping duration of less than 15 minutes. Pinpointing single muscle representations, in contrast to an overall cortical probability map, requires between 190 (FDI) and 380 (ADM) pulses on average with significant inter-subject variability. Information about the convergence properties enables the quantification of the overall quality of a mapping. Specifically, the normalized root mean square deviation between all realized stimulations versus all but one stimulations (*n* - 1) should be as low as 2% to assert valid mapping results.

Here, we applied the proposed localization method in a domain that is characterized by a clear relationship between a single cortical site and the modulated behavioral effect. Due to its genericity, the method may be adapted to domains outside the motor cortex. Cognitive brain functions tend to feature a higher trial-by-trial variability and would thus require more stimulations. In addition, complex structure-function relationships for cognitive functions often rely on distributed networks of interacting regions. Therefore, methodological extensions, such as multivariate regression approaches, are required to identify interdependent relationships between multiple neuronal populations.

Our approach was designed to increase the variance between electric field distributions via the use of random coil positions and orientations. This partially solves the problem of a missing criterion to define coil configurations a-priori [1], increases the potential information gain per pulse, and thus substantially reduces the required number of stimulations for a robust localization. Despite this efficacy, a further reduction of the number of stimulations might be feasible with optimization procedures, which identify optimal combinations of coil positions and orientations a priori. Allowing random coil positions and orientations instead of acquiring entire input-output curves for predefined coil positions/orientations establishes a highly efficient and robust experimental protocol.

In conclusion, we propose an efficient and easy to implement high-resolution TMS localization method, applicable with standard TMS hardware. This framework enables the mapping of causal structure-function relationships with a precision comparable to high-resolution neuroimaging techniques. Most importantly, the achieved mapping quality can be quantified, either online or post-experimentally. Our framework may be easily transferred to other functional domains with a single cortical representation.

## Acknowledgments

This work was partially supported by the German Science Foundation (DFG) (HA 6314/3-1 and HA 6314/9-1 to GH, KN 588/10-1 to TRK and WE 59851/2 to KW); The Lundbeck Foundation (grants no. R244-2017-196 and R313-2019-622 to AT), the NVIDIA Corporation (donation of two Titan Xp graphics cards to GH and KW). GH is supported by the Max Planck Society.

## References

[1] Weise K, Numssen O, Thielscher A, Hartwigsen G, Knösche TR. A novel approach to localize cortical TMS effects. Neuroimage 2020;209:116486. https://doi.org/10.1016/j.neuroimage.2019.116486.

[2] Gomez LJ, Dannhauer M, Peterchev AV. Fast computational optimization of TMS coil placement for individualized electric field targeting. Neuroimage 2021;228:117696. https://doi.org/10.1016/j.neuroimage.2020.117696.

[3] Classen J, Knorr U, Werhahn KJ, Schlaug G, Kunesch E, Cohen LG, et al. Multimodal output mapping of human central motor representation on different spatial scales. J Physiol 1998;512 (Pt 1):163–79. https://doi.org/10.1111/j.1469-7793.1998.163bf.x.

[4] Siebner HR. Does TMS of the precentral motor hand knob primarily stimulate the dorsal premotor cortex or the primary motor hand area? Brain Stimul 2020;13:517–8. https://doi.org/10.1016/j.brs.2019.12.015.

[5] Dubbioso R, Madsen KH, Thielscher A, Siebner HR. Multimodal finger-printing of the human precentral cortex forming the motor hand knob. Cold Spring Harbor Laboratory 2020:2020.02.11.942771. https://doi.org/10.1101/2020.02.11.942771.

[6] Fox PT, Narayana S, Tandon N, Sandoval H, Fox SP, Kochunov P, et al. Column-based model of electric field excitation of cerebral cortex. Hum Brain Mapp 2004;22:1–14. https://doi.org/10.1002/hbm.20006.

[7] Krieg TD, Salinas FS, Narayana S, Fox PT, Mogul DJ. PET-based confirmation of orientation sensitivity of TMS-induced cortical activation in humans. Brain Stimul 2013;6:898–904. https://doi.org/10.1016/j.brs.2013.05.007.

[8] Teitti S, Määttä S, Säisänen L, Könönen M, Vanninen R, Hannula H, et al. Non-primary motor areas in the human frontal lobe are connected directly to hand muscles. Neuroimage 2008;40:1243–50. https://doi.org/10.1016/j.neuroimage.2007.12.065.

[9] Diekhoff S, Uludağ K, Sparing R, Tittgemeyer M, Cavuşoğlu M, von Cramon DY, et al. Functional localization in the human brain: Gradient-Echo, Spin-Echo, and arterial spin-labeling fMRI compared with neuronavigated TMS. Hum Brain Mapp 2011;32:341–57. https://doi.org/10.1002/hbm.21024.

[10] Sarfeld A-S, Diekhoff S, Wang LE, Liuzzi G, Uludağ K, Eickhoff SB, et al. Convergence of human brain mapping tools: neuronavigated TMS parameters and fMRI activity in the hand motor area. Hum Brain Mapp 2012;33:1107–23. https://doi.org/10.1002/hbm.21272.

[11] Ahdab R, Ayache SS, Farhat WH, Mylius V, Schmidt S, Brugières P, et al. Reappraisal of the anatomical landmarks of motor and premotor cortical regions for image-guided brain navigation in TMS practice. Human Brain Mapping 2014;35:2435–47. https://doi.org/10.1002/hbm.22339.

[12] Ahdab R, Ayache SS, Brugières P, Farhat WH, Lefaucheur J-P. The Hand Motor Hotspot is not Always Located in the Hand Knob: A Neuronavigated Transcranial Magnetic Stimulation Study. Brain Topography 2016;29:590–7. https://doi.org/10.1007/s10548-016-0486-2.

[13] Vaalto S, Säisänen L, Könönen M, Julkunen P, Hukkanen T, Määttä S, et al. Corticospinal output and cortical excitation-inhibition balance in distal hand muscle representations in nonprimary motor area. Human Brain Mapping 2011;32:1692–703. https://doi.org/10.1002/hbm.21137.

[14] Neggers SFW, Langerak TR, Schutter DJLG, Mandl RCW, Ramsey NF, Lemmens PJJ, et al. A stereotactic method for image-guided transcranial magnetic stimulation validated with fMRI and motor-evoked potentials. Neuroimage 2004;21:1805–17. https://doi.org/10.1016/j.neuroimage.2003.12.006.

[15] Kleim JA, Kleim ED, Cramer SC. Systematic assessment of training-induced changes in corticospinal output to hand using frameless stereotaxic transcranial magnetic stimulation. Nat Protoc 2007;2:1675–84. https://doi.org/10.1038/nprot.2007.206.

[16] Ngomo S, Leonard G, Moffet H, Mercier C. Comparison of transcranial magnetic stimulation measures obtained at rest and under active conditions and their reliability. J Neurosci Methods 2012;205:65–71. https://doi.org/10.1016/j.jneumeth.2011.12.012.

[17] van de Ruit M, Perenboom MJL, Grey MJ. TMS brain mapping in less than two minutes. Brain Stimul 2015;8:231–9. https://doi.org/10.1016/j.brs.2014.10.020.

[18] Veldema J, Bösl K, Nowak DA. Motor Recovery of the Affected Hand in Subacute Stroke Correlates with Changes of Contralesional Cortical Hand Motor Representation. Neural Plast 2017;2017:6171903. https://doi.org/10.1155/2017/6171903.

[19] Tervo AE, Metsomaa J, Nieminen JO, Sarvas J, Ilmoniemi RJ. Automated search of stimulation targets with closed-loop transcranial magnetic stimulation. Neuroimage 2020;220:117082. https://doi.org/10.1016/j.neuroimage.2020.117082.

[20] Wassermann EM, Wang B, Zeffiro TA, Sadato N, Pascual-Leone A, Toro C, et al. Locating the motor cortex on the MRI with transcranial magnetic stimulation and PET. Neuroimage 1996;3:1–9. https://doi.org/10.1006/nimg.1996.0001.

[21] Opitz A, Legon W, Rowlands A, Bickel WK, Paulus W, Tyler WJ. Physiological observations validate finite element models for estimating subject-specific electric field distributions induced by transcranial magnetic stimulation of the human motor cortex. Neuroimage 2013;81:253–64. https://doi.org/10.1016/j.neuroimage.2013.04.067.

[22] Aonuma S, Gomez-Tames J, Laakso I, Hirata A, Takakura T, Tamura M, et al. A high-resolution computational localization method for transcranial magnetic stimulation mapping. Neuroimage 2018;172:85–93. https://doi.org/10.1016/j.neuroimage.2018.01.039.

[23] Bungert A, Antunes A, Espenhahn S, Thielscher A. Where does TMS Stimulate the Motor Cortex? Combining Electrophysiological Measurements and Realistic Field Estimates to Reveal the Affected Cortex Position. Cereb Cortex 2017;27:5083–94. https://doi.org/10.1093/cercor/bhw292.

[24] Laakso I, Murakami T, Hirata A, Ugawa Y. Where and what TMS activates: Experiments and modeling. Brain Stimul 2018;11:166–74. https://doi.org/10.1016/j.brs.2017.09.011.

[25] Geyer S, Ledberg A, Schleicher A, Kinomura S, Schormann T, Bürgel U, et al. Two different areas within the primary motor cortex of man. Nature 1996;382:805–7. https://doi.org/10.1038/382805a0.

[26] Rossi S, Antal A, Bestmann S, Bikson M, Brewer C, Brockmöller J, et al. Safety and recommendations for TMS use in healthy subjects and patient populations, with updates on training, ethical and regulatory issues: Expert Guidelines. Clin Neurophysiol 2021;132:269–306. https://doi.org/10.1016/j.clinph.2020.10.003.

[27] Mayka MA, Corcos DM, Leurgans SE, Vaillancourt DE. Three-dimensional locations and boundaries of motor and premotor cortices as defined by functional brain imaging: A meta-analysis. NeuroImage 2006;31:1453–74. https://doi.org/10.1016/j.neuroimage.2006.02.004.

[28] Yousry T. Localization of the motor hand area to a knob on the precentral gyrus. A new landmark. Brain 1997;120:141–57. https://doi.org/10.1093/brain/120.1.141.

[29] Rothwell JC, Hallett M, Berardelli A, Eisen A, Rossini P, Paulus W. Magnetic stimulation: motor evoked potentials. The International Federation of Clinical Neurophysiology. Electroencephalogr Clin Neurophysiol Suppl 1999;52:97–103.

[30] Saturnino GB, Madsen KH, Thielscher A. Electric field simulations for transcranial brain stimulation using FEM: an efficient implementation and error analysis. J Neural Eng 2019;16:066032. https://doi.org/10.1088/1741-2552/ab41ba.

[31] Thielscher A, Antunes A, Saturnino GB. Field modeling for transcranial magnetic stimulation: A useful tool to understand the physiological effects of TMS? 2015 37th Annual International Conference of the IEEE Engineering in Medicine and Biology Society (EMBC) 2015. https://doi.org/10.1109/embc.2015.7318340.

[32] Güllmar D, Haueisen J, Reichenbach JR. Influence of anisotropic electrical conductivity in white matter tissue on the EEG/MEG forward and inverse solution. A high-resolution whole head simulation study. NeuroImage 2010;51:145–63. https://doi.org/10.1016/j.neuroimage.2010.02.014.

[33] Nielsen JD, Madsen KH, Puonti O, Siebner HR, Bauer C, Madsen CG, et al. Automatic skull segmentation from MR images for realistic volume conductor models of the head: Assessment of the state-of-the-art. Neuroimage 2018;174:587–98. https://doi.org/10.1016/j.neuroimage.2018.03.001.

[34] Penny WD, Friston KJ, Ashburner JT, Kiebel SJ, Nichols TE. Statistical Parametric Mapping: The Analysis of Functional Brain Images. Elsevier; 2011.

[35] Gaser C, Dahnke R, Kurth F, Luders E. A Computational Anatomy Toolbox for the Analysis of Structural MRI Data. NeuroImage n.d.

[36] Wagner T, Gangitano M, Romero R, Théoret H, Kobayashi M, Anschel D, et al. Intracranial measurement of current densities induced by transcranial magnetic stimulation in the human brain. Neurosci Lett 2004;354:91–4. https://doi.org/10.1016/s0304-3940(03)00861-9.

[37] Fischl B, Dale AM, Sereno MI, Tootell RBH, Rosen BR. A Coordinate System for the Cortical Surface. NeuroImage 1998;7:S740. https://doi.org/10.1016/s1053-8119(18)31573-8.

[38] Dale AM, Fischl B, Sereno MI. Cortical Surface-Based Analysis. NeuroImage 1999;9:179–94. https://doi.org/10.1006/nimg.1998.0395.

[39] Glasser MF, Coalson TS, Robinson EC, Hacker CD, Harwell J, Yacoub E, et al. A multi-modal parcellation of human cerebral cortex. Nature 2016;536:171–8. https://doi.org/10.1038/nature18933.

[40] Aberra AS, Wang B, Grill WM, Peterchev AV. Simulation of transcranial magnetic stimulation in head model with morphologically-realistic cortical neurons. Brain Stimul 2020;13:175–89. https://doi.org/10.1016/j.brs.2019.10.002.

[41] Alavi SMM, Goetz SM, Peterchev AV. Optimal Estimation of Neural Recruitment Curves Using Fisher Information: Application to Transcranial Magnetic Stimulation. IEEE Transactions on Neural Systems and Rehabilitation Engineering 2019;27:1320–30. https://doi.org/10.1109/tnsre.2019.2914475.

[42] Goetz SM, Luber B, Lisanby SH, Peterchev AV. A Novel Model Incorporating Two Variability Sources for Describing Motor Evoked Potentials. Brain Stimulation 2014;7:541–52. https://doi.org/10.1016/j.brs.2014.03.002.

[43] Goetz SM, Mahdi Alavi SM, Deng Z-D, Peterchev AV. Statistical Model of Motor-Evoked Potentials. IEEE Transactions on Neural Systems and Rehabilitation Engineering 2019;27:1539–45. https://doi.org/10.1109/tnsre.2019.2926543.

[44] Peterchev AV, Goetz SM, Westin GG, Luber B, Lisanby SH. Pulse width dependence of motor threshold and input–output curve characterized with controllable pulse parameter transcranial magnetic stimulation. Clinical Neurophysiology 2013;124:1364–72. https://doi.org/10.1016/j.clinph.2013.01.011.

[45] Nielsen JF. Logarithmic Distribution of Amplitudes of Compound Muscle Action Potentials Evoked by Transcranial Magnetic Stimulation. Journal of Clinical Neurophysiology 1996;13:423–34. https://doi.org/10.1097/00004691-199609000-00005.

[46] Dum R, Strick P. Motor areas in the frontal lobe of the primate. Physiology & Behavior 2002;77:677–82. https://doi.org/10.1016/s0031-9384(02)00929-0.

[47] Ninomiya T, Inoue K-I, Hoshi E, Takada M. Layer specificity of inputs from supplementary motor area and dorsal premotor cortex to primary motor cortex in macaque monkeys. Scientific Reports 2019;9. https://doi.org/10.1038/s41598-019-54220-z.

[48] Schaworonkow N, Gordon PC, Belardinelli P, Ziemann U, Bergmann TO, Zrenner C. μ-Rhythm Extracted With Personalized EEG Filters Correlates With Corticospinal Excitability in Real-Time Phase-Triggered EEG-TMS. Frontiers in Neuroscience 2018;12. https://doi.org/10.3389/fnins.2018.00954.

[49] Zrenner C, Desideri D, Belardinelli P, Ziemann U. Real-time EEG-defined excitability states determine efficacy of TMS-induced plasticity in human motor cortex. Brain Stimul 2018;11:374–89. https://doi.org/10.1016/j.brs.2017.11.016.

[50] Bergmann TO, Lieb A, Zrenner C, Ziemann U. Pulsed Facilitation of Corticospinal Excitability by the Sensorimotor μ-Alpha Rhythm. The Journal of Neuroscience 2019;39:10034–43. https://doi.org/10.1523/jneurosci.1730-19.2019.

[51] Triesch J, Zrenner C, Ziemann U. Modeling TMS-induced I-waves in human motor cortex. Progress in Brain Research 2015:105–24. https://doi.org/10.1016/bs.pbr.2015.07.001.

[52] Huber L, Finn ES, Handwerker DA, Bönstrup M, Glen DR, Kashyap S, et al. Sub-millimeter fMRI reveals multiple topographical digit representations that form action maps in human motor cortex. NeuroImage 2020;208:116463. https://doi.org/10.1016/j.neuroimage.2019.116463.

[53] Wilson SA, Thickbroom GW, Mastaglia FL. Transcranial magnetic stimulation mapping of the motor cortex in normal subjects. Journal of the Neurological Sciences 1993;118:134–44. https://doi.org/10.1016/0022-510x(93)90102-5.

[54] Martuzzi R, van der Zwaag W, Farthouat J, Gruetter R, Blanke O. Human finger somatotopy in areas 3b, 1, and 2: a 7T fMRI study using a natural stimulus. Hum Brain Mapp 2014;35:213–26. https://doi.org/10.1002/hbm.22172.

[55] Raffin E, Pellegrino G, Di Lazzaro V, Thielscher A, Siebner HR. Bringing transcranial mapping into shape: Sulcus-aligned mapping captures motor somatotopy in human primary motor hand area. Neuroimage 2015;120:164–75. https://doi.org/10.1016/j.neuroimage.2015.07.024.

[56] Raffin E, Siebner HR. Use-Dependent Plasticity in Human Primary Motor Hand Area: Synergistic Interplay Between Training and Immobilization. Cerebral Cortex 2019;29:356–71. https://doi.org/10.1093/cercor/bhy226.

[57] Dubbioso R, Raffin E, Karabanov A, Thielscher A, Siebner HR. Centre-surround organization of fast sensorimotor integration in human motor hand area. Neuroimage 2017;158:37–47. https://doi.org/10.1016/j.neuroimage.2017.06.063.

[58] Goldsworthy MR, Hordacre B, Ridding MC. Minimum number of trials required for within- and between-session reliability of TMS measures of corticospinal excitability. Neuroscience 2016;320:205–9. https://doi.org/10.1016/j.neuroscience.2016.02.012.

