## Supplemental Material for "Efficient high-resolution TMS mapping of the human motor cortex by nonlinear regression" (from another version of this preprint: version 1, 2021-03-12)

Ole Numssen[1]*, Anna-Leah Zier[1,4], Axel Thielscher[2,3], Gesa Hartwigsen[1], Thomas R. Knösche[4,5], Konstantin Weise[4,6]

[1]*Lise Meitner Research Group "Cognition and Plasticity", Max Planck Institute for Human Cognitive and Brain Sciences, Stephanstr. 1a, 04103 Leipzig, Germany.*

[2]*Danish Research Centre for Magnetic Resonance, Centre for Functional and Diagnostic Imaging and Research, Copenhagen University Hospital Amager and Hvidovre, Denmark.*

[3]*Technical University of Denmark, Center for Magnetic Resonance, Department of Health Technology, Kongens Lyngby, Denmark.*

[4]*Methods and Development Group "Brain Networks", Max Planck Institute for Human Cognitive and Brain Sciences, Stephanstr. 1a, 04103 Leipzig, Germany.*

[5]*Technische Universität Ilmenau, Institute of Biomedical Engineering and Informatics, Gustav-Kirchhoff-Straße 2, 98693 Ilmenau, Germany.*

[6]*Technische Universität Ilmenau, Advanced Electromagnetics Group, Helmholtzplatz 2, 98693 Ilmenau, Germany.*

* CORRESPONDING AUTHOR

*Ole Numssen; Max Planck Institute for Human Cognitive and Brain Sciences, Stephanstr. 1a, 04103 Leipzig, Germany;, phone: +49 341 9940-2010*

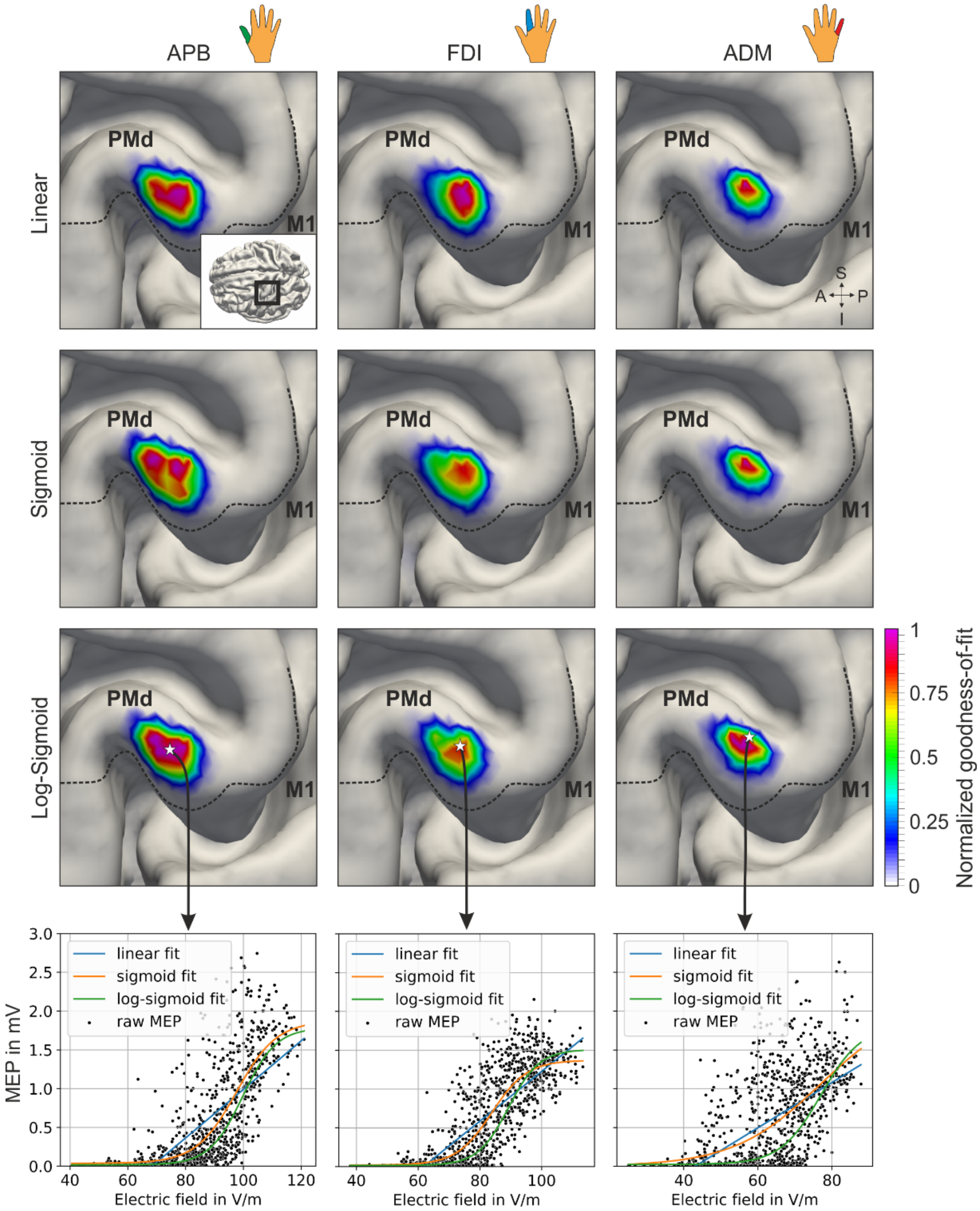


**Figure S1:** Comparison between the different curve fitting procedures for subject 12. The first three columns show the normalized coefficients of determination ($R^2$) for APB, FDI, and ADM, respectively, considering the tangential component of the electric field. The color scale is the same as in Fig. 3. The last column highlights the identified hotspots for the three digits from the magnitude of the electric field. The boundary between dorsal premotor cortex PMd/BA6 and the primary motor cortex M1/BA4 are determined with the HCP-MMP atlas (Glasser et al. 2016). $R^2$ values were potentiated by a factor of 20 for the sake of visualization.

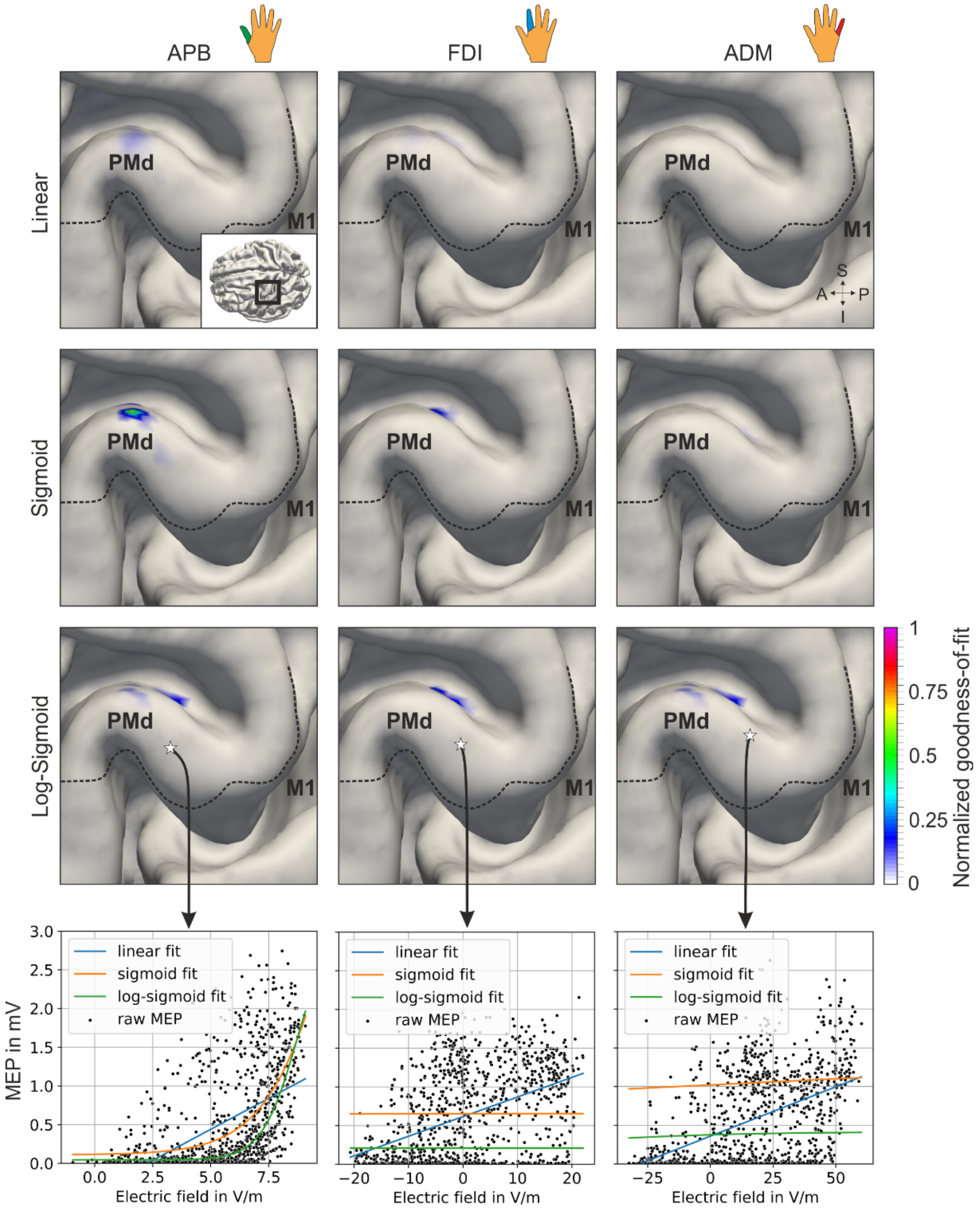


**Figure S2:** Comparison between the different curve fitting procedures for subject 12. The first three columns show the normalized coefficients of determination ($R^2$) for APB, FDI, and ADM, respectively, considering the normal component of the electric field. The color scale is the same as in Fig. 3. The last column highlights the identified hotspots for the three digits from the magnitude of the electric field. The boundary between dorsal premotor cortex PMd/BA6 and the primary motor cortex M1/BA4 are determined with the HCP-MMP atlas (Glasser et al. 2016). $R^2$ values were potentiated by a factor of 20 for the sake of visualization.

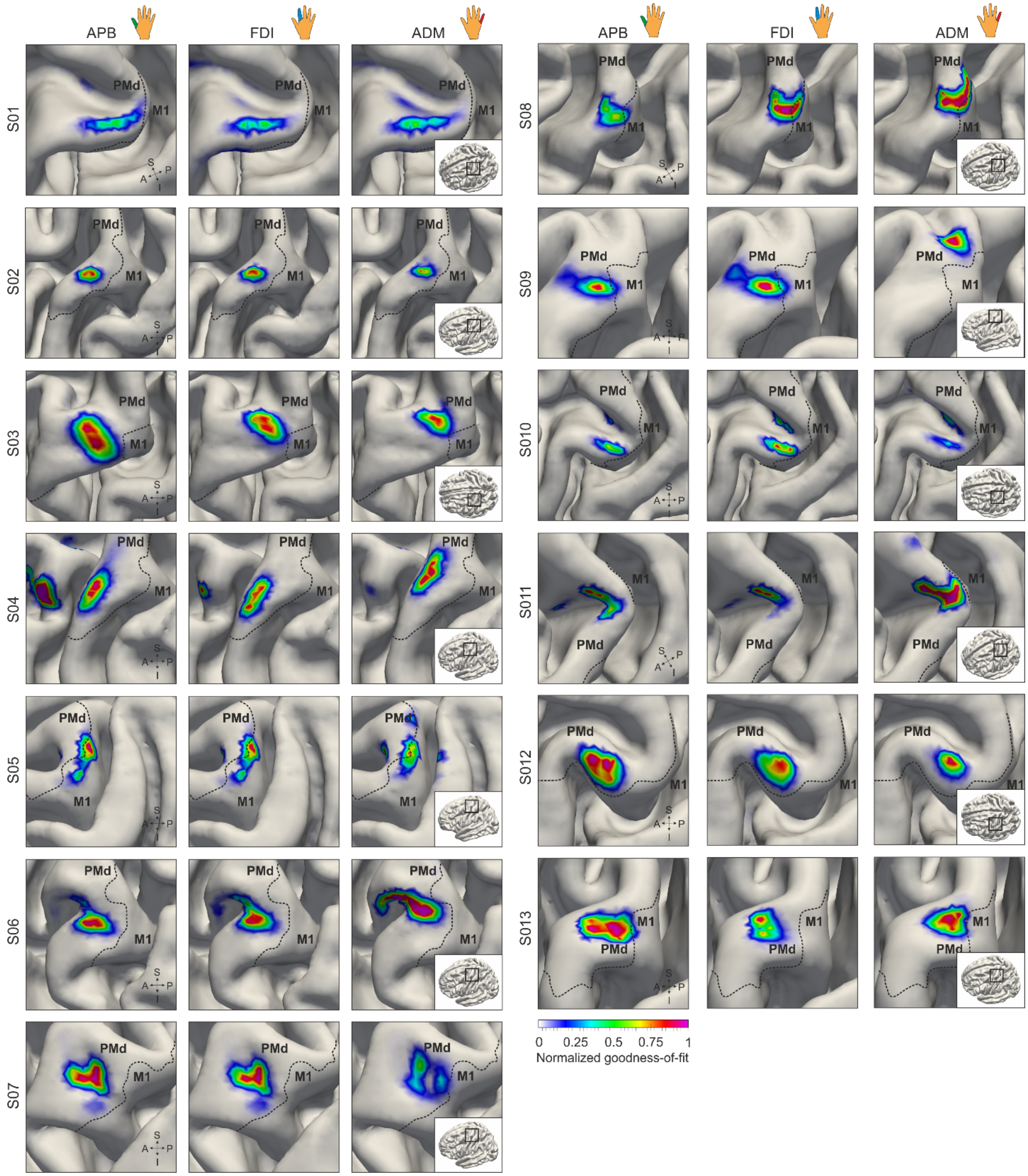


**Figure S3:** Localization results from all 13 subjects using nonlinear regression with a sigmoidal function. The first three columns show the normalized coefficients of determination ($R^2$) for APB, FDI, and ADM, respectively, considering the tangential component of the electric field. The color scale is the same as in Fig. 4. The first subject S01 was measured with pseudo-random stimulation intensities, instead of the fixed intensity that was used for subsequent subjects. The boundary between dorsal premotor cortex PMd/BA6 and the primary motor cortex M1/BA4 are determined with the HCP-MMP atlas (Glasser et al. 2016). $R^2$ values were potentiated by a factor of 20 for the sake of visualization.

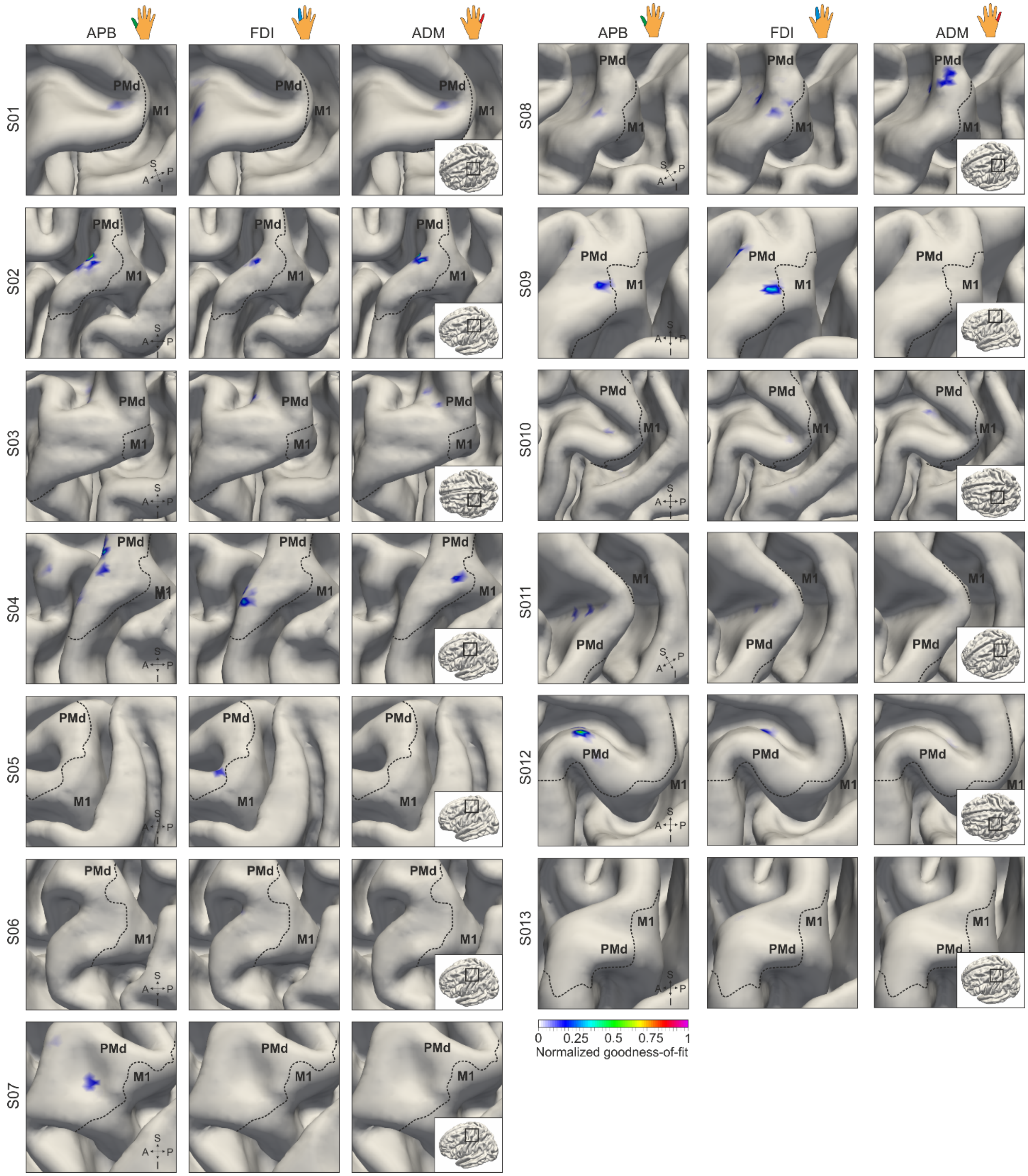


**Figure S4:** Localization results from all 13 subjects using nonlinear regression with a sigmoidal function. The first three columns show the normalized coefficients of determination ($R^2$) for APB, FDI, and ADM, respectively, considering the normal component of the electric field. The color scale is the same as in Fig. 4. The first subject S01 was measured with pseudo-random stimulation intensities, instead of the fixed intensity that was used for subsequent subjects. The boundary between dorsal premotor cortex PMd/BA6 and the primary motor cortex M1/BA4 are determined with the HCP-MMP atlas (Glasser et al. 2016). $R^2$ values were potentiated by a factor of 20 for the sake of visualization.

**Table S1:** Maximum $R^2$ values for different fit functions

| | APB | | | FDI | | | ADM | | |
|---|---|---|---|---|---|---|---|---|---|
| Subject | linear | sigmoid | log sigmoid | linear | sigmoid | log sigmoid | linear | sigmoid | log sigmoid |
| S01 | 0.50 | 0.70 | 0.69 | 0.58 | 0.71 | 0.70 | 0.46 | 0.75 | 0.74 |
| S02 | 0.31 | 0.38 | 0.36 | 0.44 | 0.56 | 0.54 | 0.35 | 0.49 | 0.47 |
| S03 | 0.33 | 0.35 | 0.33 | 0.66 | 0.7 | 0.69 | 0.51 | 0.55 | 0.54 |
| S04 | 0.32 | 0.34 | 0.28 | 0.39 | 0.52 | 0.47 | 0.35 | 0.47 | 0.43 |
| S05 | 0.32 | 0.55 | 0.51 | 0.37 | 0.63 | 0.57 | 0.29 | 0.49 | 0.44 |
| S06 | 0.55 | 0.69 | 0.68 | 0.64 | 0.76 | 0.75 | 0.46 | 0.58 | 0.56 |
| S07 | 0.42 | 0.57 | 0.54 | 0.51 | 0.63 | 0.60 | 0.5 | 0.69 | 0.67 |
| S08 | 0.41 | 0.58 | 0.55 | 0.51 | 0.70 | 0.69 | 0.24 | 0.44 | 0.41 |
| S09 | 0.31 | 0.35 | 0.33 | 0.42 | 0.47 | 0.44 | 0.27 | 0.30 | 0.28 |
| S10 | 0.41 | 0.58 | 0.51 | 0.42 | 0.59 | 0.49 | 0.37 | 0.59 | 0.50 |
| S11 | 0.31 | 0.36 | 0.33 | 0.38 | 0.42 | 0.39 | 0.27 | 0.29 | 0.26 |
| S12 | 0.52 | 0.61 | 0.60 | 0.62 | 0.67 | 0.65 | 0.58 | 0.64 | 0.62 |
| S13 | 0.15 | 0.20 | 0.13 | 0.15 | 0.24 | 0.15 | 0.19 | 0.33 | 0.27 |

*Notes:* For each mapping solution per subject, muscle, and fit function the maximum $R^2$ values are shown.

**Table S2:** Resting motor thresholds determined in the validation study for the different coil positions

| Subject | Target | Coil positions | | | | | | |
|---|---|---|---|---|---|---|---|---|
| | | **opt** | **opt$_{-45°}$** | **opt$_{45°}$** | **pos** | **ant** | **inf** | **sup** |
| S03 | FDI | **42** | 47 | 46 | 44 | 41 | 43 | 43 |
| | ADM | **40** | 53 | 46 | 44 | 44 | 43 | 43 |
| | APB | **42** | 53 | **42** | 44 | 44 | 45 | 43 |
| S04 | FDI | 36 | 49 | 39 | 39 | 40 | 41 | **35** |
| | ADM | **36** | 46 | 40 | 38 | 37 | 39 | 39 |
| | APB | **36** | 43 | 37 | **36** | **36** | 37 | **36** |
| S05 | FDI | **48** | 52 | 64 | 52 | 57 | 61 | **48** |
| | ADM | 48 | 59 | 54 | 49 | 58 | 58 | **47** |
| | APB | 49 | 56 | 57 | 46 | **45** | 50 | 46 |
| S08 | FDI | **52** | 64 | 60 | 57 | 55 | 60 | 60 |
| | ADM | 59 | **58** | 68 | **58** | 62 | 62 | 62 |
| | APB | **55** | 66 | 62 | 60 | 59 | 59 | 58 |
| S09 | FDI | **47** | 53 | 54 | 51 | 49 | 51 | 50 |
| | ADM | 55 | 54 | 62 | 60 | **51** | 55 | 62 |
| | APB | 48 | 58 | **45** | 55 | **45** | 49 | 52 |
| S10 | FDI | **42** | 50 | 50 | 51 | 43 | 46 | 47 |
| | ADM | **43** | 48 | 54 | 49 | 44 | 46 | 49 |
| | APB | **40** | 50 | 47 | 48 | 43 | 44 | 47 |
| S11 | FDI | **72** | 90 | 87 | 72 | 88 | 89 | 74 |
| | ADM | **72** | 84 | 88 | 74 | 86 | 89 | 72 |
| | APB | **72** | 96 | 78 | 73 | 80 | 84 | 75 |
| S12 | FDI | **46** | 56 | 60 | 51 | 48 | 50 | **46** |
| | ADM | 47 | 59 | 58 | 52 | **46** | 48 | 49 |
| | APB | **50** | 64 | 51 | 51 | 57 | 54 | **50** |

*Notes:* opt: optimal coil position including ±45° orientations; pos, ant, inf and sup denote posterior, anterior, inferior and superior coil positions, respectively, shifted by 7.5 mm with respect to the optimal coil position. Minimum values highlighted in bold.
